# MicroRNA hsa-miR-203a-3p promotes H1N1 and NDV Virus Replication by suppressing the IFNA signaling

**DOI:** 10.1101/2024.04.30.591875

**Authors:** Pramod kumar, Ashish Kumar, Akhilesh Kumar, Himanshu Kumar

## Abstract

MicroRNAs (miRNAs) are small non-coding molecules that act as essential post-transcriptional regulators in various biological processes. Many studies suggest that miRNAs may modulate the host’s immune response during virus infections. We analyzed publicly available transcriptomics data involving infection with different RNA viruses and selected the most prominent candidate, miR-203a-3p. miR-203a-3p is upregulated during H7N1, HCV, and HIV+HPV infections. Interestingly, pathway analysis of microRNA-targeted genes shows that miR-203a-3p targets the type–I interferon pathway. In this study, we report that the expression of miR-203a-3p is elevated in response to polyinosinic-polycytidylic acid [poly(I:C)] transfection and infection with RNA viruses like Newcastle Disease Virus (NDV) and H1N1 influenza virus. We found that overexpression of miR-203-3p promotes the replication of H1N1 virus by suppressing the host’s type-I interferons and interferon-stimulated genes. In addition, miR-203a-3p overexpression reduced the expression of ISGs, and is attributed to the binding of miR-203a-3p to 3’ UTRs of Janus-activated kinase 1 (JAK1), STAT1, SOCS3, and multiple IFNA transcripts, as shown by luciferase and Ago-2 pulldown assays. Altogether, these findings strongly suggest that miR-203a-3p acts as a pro-viral molecule during H1N1 and NDV infection by targeting the host’s IFN signaling pathways.

## Introduction

RNA viruses like influenza, SARS, MERS, and SARS-CoV-2 have been the cause of several pandemics and epidemics in recent past and therefore pose a significant threat to global human health. These viruses are tackled by the host’s innate immune system, which is also the first line of defense against invading pathogens. The innate immune system consists of specific pattern recognition receptors (PRRs)(1–3) that sense the conserved molecular signatures associated with viruses such as viral protein, RNA with 5′-triphosphate ends, viral DNA, single-stranded RNA (ssRNA), and double-stranded RNA (dsRNA). PRRs activate a plethora of downstream signaling culminating in the activation of IFN regulatory factor (IRFs) and nuclear factor-kB (NF-kB), which act as transcription factors for Type-I and III interferons and pro-inflammatory cytokines. (4–6) While interferons work in an autocrine and paracrine manner to help nearby cells attain an antiviral state, the pro-inflammatory cytokines invite different immune cells to the site of infection. (3)Type-I interferons have a very prominent antiviral effect because they induce an array of genes called interferon-stimulated genes (ISGs) that exert a pleiotropic effect on viral replication. (7) The effector protein of IFN signaling pathways like STAT (signal transducer and activator of transcription) complexes activates and regulates the expression of target genes (8, 9). Where STAT1-STAT2-IRF9 complex acts as a transcription factor for interferon sensitive response element 9 (ISRE9). Itinitiates the transcription of several antiviral ISGs like, GBPs, IFITMs, TRIMs, NOS2, IRFs, STATs, IFNAs, OASs, MXs, ISG15, and viperin(10, 11). This complex process of antiviral signaling is carefully regulated by post-transcriptional and post-translational mechanisms. While post-translational mechanisms, such as ubiquitination or phosphorylation of IFNs, IRFs, JAKs, and STATs, regulate the activation or degradation of these proteins, post-transcriptional regulators like microRNAs (miRNAs) also plays a crucial role in fine-tuning antiviral immune responses (12) miRNAs are single-stranded non-coding RNAs (ncRNAs) ranging between 18-21 nucleotides. miRNAs help regulate the gene expression by binding to the 3’UTR (3’ untranslated region) region of target transcripts and repressing their translation. Transcriptomic studies have shown that the expression of many miRNAs is induced upon RNA virus infection. These miRNAs then target innate immune signaling pathways and modulate the antiviral response(13). miRNAs can also bind to the genomic RNA of viruses, directly affecting their transcription and viral replication(14, 15).

In this study, we have identified miR-203a-3p, which is induced upon RNA virus infection and poly(I:C) transfection. miR-203a-3p targets the 3’UTR of the IFNA pathway genes (IFNAR1, IFNA2,4,7,10,14,16,17,21). We confirmed that miR-203a-3p forms a complex with Argonaut 2 (Ago2) protein, subsequently recruiting it to the transcripts of IFNA signaling pathway genes, including JAK1 and STAT1. Ectopic expression of miR-203a-3p in HEK-293T, A549, and HeLa cells diminishes the type-I interferon mediated antiviral response, leading to an increase in viral load.

Upon infection viral RNA is sensed by Toll-like receptor 3 (TLR3), TLR7, and TLR8 located on the endosomal membrane and RIG-I and MDA5 in the cytosol inducing type-I interferons necessary to mount an effective antiviral immune response.

The type-I interferons (specially IFNα/β) are well characterized for the primary innate immune response and are necessary for defending against viruses (16–18). Type-I interferons are a family of cytokines dedicated to immunity against viruses, bacteria, parasites, and fungi. In humans and mice a large number of cells ( T-cells, B-cells, NK cells, endothelial cells, macrophage, fibroblasts, and other cells express receptors for the type-I interferons on their surface(19, 20). IFN receptors are transmembrane receptors made up of two subunits IFNAR1 and IFNAR2. IFNAR forms a dimer upon binding of interferon (IFNα/β), dimerization in turn activates JAK1 and tyrosine kinase 2 (TYK2) resulting in activation of STAT1 and STAT2 which forms an IFN-stimulated gene factor 3 (ISGF3). Not only does STAT1/2 forms dimer but STAT1/STAT1 also forms a homodimer also known as IFNγ activating factor (GAF), where ISGF3 and GAF bind to the ISREs and IFNγ activation sites leading to activation of more than 100 ISGs, whose integrated action results in an antiviral state(21, 22). It is also reported not only STAT1/STAT1 homodimer and STAT1/STAT2 heterodimer but STAT4 can also mediate the downstream signaling in absence of STAT1 and results in the activation of many antiviral genes(23, 24). In humans, plasmacytoid dendritic cells (PDCs) are uniquely specialized for secretion of high levels of type I interferons; in general, all nucleated cells secret Type-I interferon but PDCs secrete high levels of type-I interferons through activation of endosomal TLR7 and TLR9. Release of Type-I interferon during viral infection triggers a signaling impulse that results in a expression of ISGs which have direct and indirect inhibitory effects on viral replication.

We found that miRNA-203a-3p has the ability to modulate RNA virus infection through two distinct mechanisms. It diminishes the innate immune responses in host cells by targeting type-I interferon signaling genes and ISGs and to the viral genome but does not have any significant effect on viral replication.There are no other reports found on the one miRNA targeting both IFN genes and viral genome; however, a similar study was published by our group previously, where miR-485 targeted both innate immune gene (RIG-I) of host and genomes of Newcastle disease virus (NDV) and influenza viruses (25).

## Material and Methods

NCBI-GEO database was queried for miRNA profiling experiments and databases involving H7N1, HCV, and HIV+HPV co-infection (GSE150782, GSE40744, and GSE81137) were selected. Identification of differentially expressed miRNAs in control and infected samples of the given datasets was done using GEO2R online tool (26). TargetScan 7.1(27–29), and miRabel(30, 31) were used to predict the target gens of selected miRNA.

### Cells, transfection, plasmids, mimics and inhibitors

A549 human adenocarcinomic alveolar basal epithelial cells (Cell Repository, NCCS, India), HeLa cervical cancer cells (Cell Repository, NCCS, India), and HEK293T human embryonic kidney cells (ATCC, CRL-3216), were cultured in Dulbecco’s modified Eagle’s medium (DMEM) supplemented with 10% fetal bovine serum (FBS) and 1% penicillin-streptomycin. For transfection 6, 12, or 24 well cell culture plates were used according to experiment requirements, when cells reached 70-90% of confluency, plasmid, mimics (Invitrogen) and negative control or poly I:C were transfected according to manufacturer protocol in HEK293T and HeLa cells using Lipofectamine 2000 (Invitrogen), whereas Lipofectamine 3000 (Invitrogen) was used for A549, and respective samples were collected at desired time points.

### Viruses and infection

Influenza virus (strain PR8, A/PR8/H1N1), GFP tagged NDV (strain LaSota) viruses were used in this study. Cells were seeded in 6 and 12 well tissue culture plates at a density of 0.3x106 and 0.1x106 cells/well. 24 hr. post-post-transfection cells were washed with SF-DMEM or 1x Phosphate-buffered saline (PBS) and infected with NDV-GFP at .5 MOI and PR8 at 0.5, 1, and 2 MOI. And virus was re-suspended in SF-DMEM and added to the cells. After 1 h, cells were washed again with SF-DMEM or 1x Phosphate-buffered saline (PBS) and incubated in complete DMEM and supplemented with 1% FBS. Replacement media for PR8 infection was additionally supplemented with L-1-Tosylamide-2-phenylethyl chloromethyl ketone (TPCK)-treated trypsin (1 μg/ml). NDV-GFP was gifted by Dr. Peter Palese, Icahn School of Medicine at Mount Sinai, New York, USA.

### Luciferase reporter assay

Cells were cultured in 24 well plates and were lysed using passive lysis buffer provided with the assay kit. 50µl of Dual-Glo Luciferase Reagent was added to 50µl of cell lysate in each well of opaque (white) 96 well plate. Luminescence of firefly luciferase was measured after mixing by gently tapping the plate. To measure the Renilla luciferase activity equal volume of Stop & Glow reagent was added per well. After 10 minutes of incubation luminescence of renilla luciferase was measured. Luminescence intensity of renilla luciferase was used to normalize the firefly luciferase luminescence.

### Quantitative real-time reverse transcription-PCR

Total RNA was extracted using TRIzol reagent (Invitrogen) and it was used to synthesize cDNA using the iScript cDNA synthesis kit (Bio-Rad) according to the manufacturer’s protocol. Gene expression measured by Quantitative real-time PCR using gene specific primers with CYBR green chemistry (Bio-Rad), 18S RNA was used as an internal control for normalization. To check miRNA expression TaqMan universal PCR master mix (Applied Biosystems) based real-time PCR was performed, where miR-203a-3p and U6 specific TaqMan assay was used. Here U6 was used as an internal control for normalization.

### Cytofluorometric analysis

Cells were seeded in six-well plates at approximate density of 60–70%. In all, 24 h later, cells were transfected with pMIR-203a-3p and pMIR-empty as a control vector. After 24 cells were washed with SF-DMEM or 1x Phosphate-buffered saline (PBS) and infected with NDV-GFP at 1.5 MOI for 1 hour. Cell monolayer was washed with PBS, and medium was replaced with 1% complete DMEM. The next day cells were detached using trypsin-EDTA solution, washed with PBS and fixed with 4% paraformaldehyde for 5 min. Cells were washed twice with PBS and resuspended in PBS for cytofluorometric analysis on a BD FACSaria II flow cytometer.

### RNA immunoprecipitation

RNA immunoprecipitation was performed as describe in the previous articles(32, 33). HEK 293T cells were lysed in lysis buffer (0.5% NP-40, 150 mM KCl, 25 mM tris-glycine (pH 7.5) and incubated with M2 Flag affinity beads overnight Thereafter, the lysate was washed 5 times with washing buffer (300 mM NaCl, 50 mM Tris-glycine [pH 7.5], 5 mM MgCl2, 0.05% NP-40). The extraction of RNA from immunoprecipitated ribonucleoproteins (RNPs) was performed using Trizol reagent according to the manufacturer’s protocol.

### Measurement of viral titer

MDCK cells were infected with virus (PR8 and NDV) in the serum-free MEM medium with the cell culture supernatant containing virus. after 1h of virus infection cells were washed with 1X PBS. For plaque assay, cells were overlaid with 1.2% Avicel (Sigma-Aldrich; equivalent to Avicel RC-581 from FMC Corp.) in 2 ml maintenance medium and incubated at 37°C with 5% CO2. After 96 h, Avicel containing medium was removed and the cells were fixed with 4% paraformaldehyde for 20 min at 4°C. Cells were stained with 0.5% crystal violet solution and plaques were counted under microscope in transmitted light. For the 50% tissue culture infective dose (TCID50) assay, cell monolayers were fixed and stained with 1% crystal violet after 96 h of infection and the TCID50 was calculated based on the Reed and Muench method as described previously(34, 35).

## Results

### miR-203a-3p is upregulated during RNA virus infection

miRNAs are key regulators of innate immune signaling and play a crucial role during RNA virus infection by directly targeting the viral genome. Therefore, we compared publically available microarray data sets (GSE150782, GSE40744, and GSE81137) of different viruses (H7N1, HCV, and HIV+HPV). Dysregulated miRNAs were selected on the basis of the p-value (< 0.05) and log2 fold change (>0.5 & < -0.5) criterions. miR-203a-3p, miR-378c and miR-199a-5p were the only miRNAs that were significantly upregulated in all three datasets. Next, these three miRNAs were analyzed for binding targets using the TargetScan 7.2 and miRabel software. The genes targeted by miR-203a-3p, miR-378c and miR-199a-5p analyzed through BioPlanet and KEGG pathway(36–39), revealed that miR-203a-3p specifically influences the regulation of Interferon alpha (IFNA) signaling, the activation of TRAF6-mediated IRF7 and the signaling pathway involving RIG-I-like receptors (Fig. S1 A-D), that play a vital role during RNA virus infections. However, miRNA-378c and miR-199a-5p does not target innate immune pathways (Fig. S2 A-D). As miR-203a-3p targets antiviral pathways and is upregulated during RNA virus infection, we decided to further study its role during RNA virus infection. We analyzed the expression of miRNA-203a-3p in HEK293, HeLa, and A549 cells and found that it is induced multifold during NDV and A/PR8/H1N1 infection (fig. 1E-H). Similarly, we observed an increase in expression of miR-203a-3p when A549, 293T, and A549 cells were transfected with poly(I:C), which simulates virus infection by activating TLR3 (Fig. S1 F).

**Figure 1:**
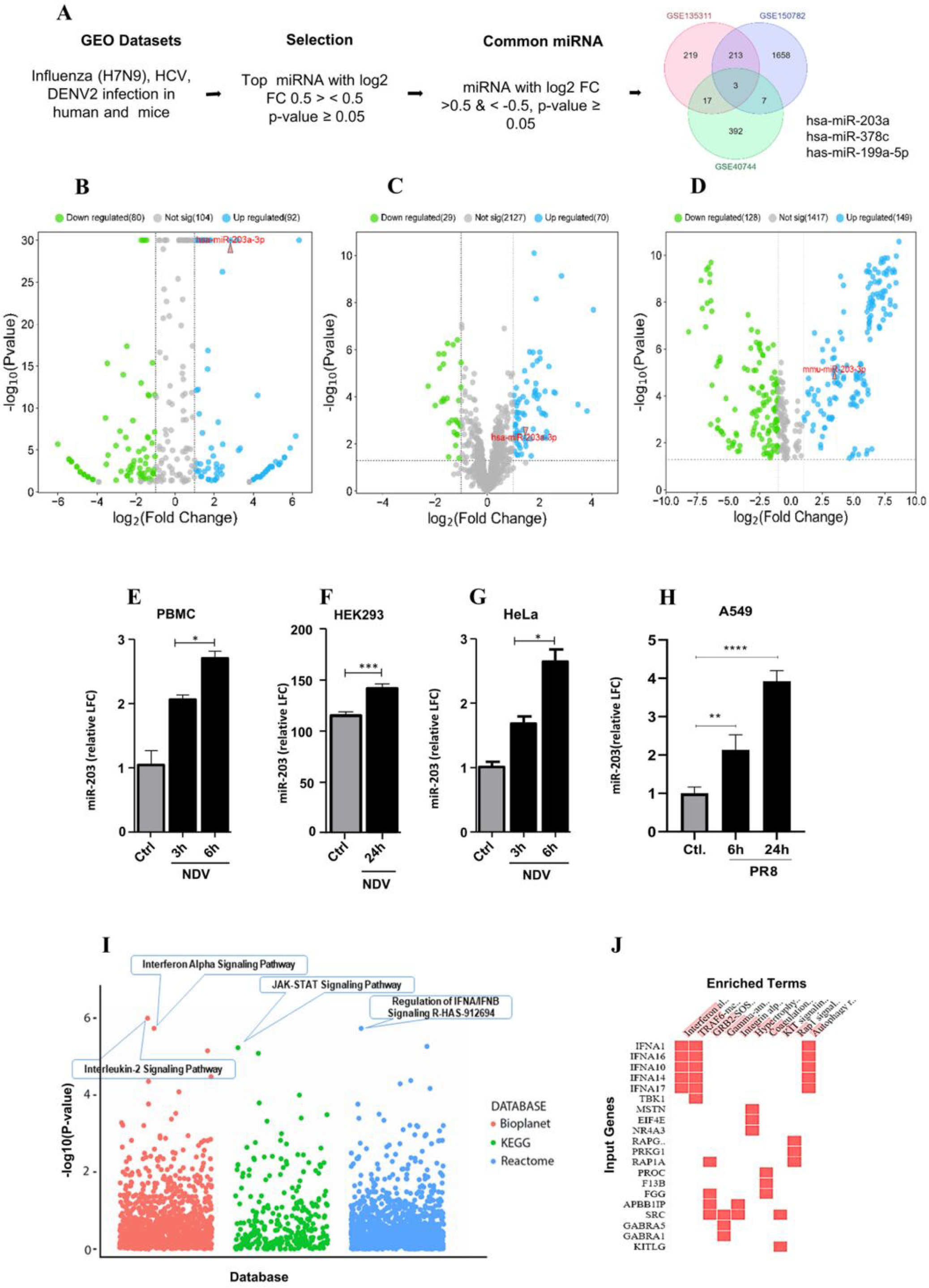
Identification and induction of miR-203-3p. (A) Different viruses H7N9, HCV, DENV2 microarray data sets (GSE150782, GSE40744, GSE135311) taken from Gene Expression Omnibus (GEO) database and reanalyzed for differential miRNAs expression using GEO2R tool. Dysregulated miRNA was further ruled out on the base of the p-value > 0.05 and log2 FC >0.5 & < -0.5. in the final output, only miR-199a-5p, miR-378c and miR-203a-3p were found to be significantly upregulated in all three datasets. (B-D) Volcano plot showing the indicating the expression of miR-203a-3p. (E-G) showing miR-203a-3p overexpression in PBMC, HEK293 and A549 cell lines upon NDV and PR8 infection at different time points. (H) miR-203a-3p overexpression upon transfection of poly I:C at 24 hours in A549 cell line. (I) pathway analysis showing the top hits for pathways in different databses Bioplanet, KEGG and Reactome, top significant pathways have been highlighted. (I) showing genes involved in top enriched pathway. The level of significance is indicated by * (p < 0.01), ** (p < 0.001) *** (p < 0.0001), **** (p < 0.00001).

### miR-203a-3p directly targets the 3’UTR of IFNA signaling regulation pathway

In-silico analysis indicate that miR-203a-3p binds to 3’UTR of IFNA subtype genes (IFNAR1, IFNA2,4,7,10,14,16,17,21), STAT1 (Fig. 2A). IFNA is a predominant member of Type-I interferon and a key contributor to the antiviral response. Upon viral infection, Type-I interferons inducesinterferon-stimulated genes (ISGs), which inhibit viral replication, and upregulate the effector function of immune cells. Therefore, we stimulated A549 cells with IFN-β and analyzed the ISRE activity by luciferase (interferon-stimulated response element) assay after overexpression of miR-203a-3p. To overexpress, the miR-203a-3p was cloned into the pMIR-REPORT vector after removing the luciferase gene, hereafter referred to as p203a. Results indicate that miR-203a-3p overexpression greatly reduces ISRE activity, indicating diminished ISG activation or type-I interferon signaling. Next, we wanted to analyze if miR-203a-3p regulates the expression of IFNAR1, IFNA1, 4, 7, and 16, JAK1, STAT1, and SOCS3 and genes. To this end, we cloned full-length UTRs of these genes downstream to the luciferase reporter gene in the pMIR-REPORT vector. HEK293T cells were co-transfected with luciferase reporter plasmids along with p203a or empty vector cells were lysed and subjected to luciferase assay after 48 hours of transfection. As expected, overexpression of miR-203a-3p greatly diminished the luciferase activity (Fig. 2B-I), indicating that miR-203a-3p binds to 3’UTR of these genes and resulting in the downregulation of their expression.To determine whether miR-203a-3p regulates the IFNA subtype genes (IFNAR1, IFNA2,4,7,10,14,16,17,21), STAT1 transcriptionally or post-transcriptionally we performed Ago2 pulldown experiment to confirm the miRNA-mRNA binding and quantify the IFNA subtype genes, and association with Ago2 protein. This protein involved in RNAi mediated gene silencing and it is a main component of RNA induced silencing complex(RISC)(40). Here we overexpressed the flag-tagged Ago2 in HEK293T cells which were later infected with NDV or A/PR8/H1N1 at an MOI of 1.5 for 24 hour. the amount of IFNA subtype transcripts that coimmunoprecipitated with miR-203a-3p were then quantified. We observed that the amount of type-I IFN genes specially IFN-β and Pan-α were in abundance in comparision of negative control(fig: 2J-K). it shows that ago2 facilitates the binding of miR-203a-3p to the type-I IFN genes IFN-β and Pan-α. We also assessed the IRF7 as a miR-203a-3p non target transcript control which shows non significant binding in both miR-203a-3p and negative control(fig: 2L).

**Figure 2:**
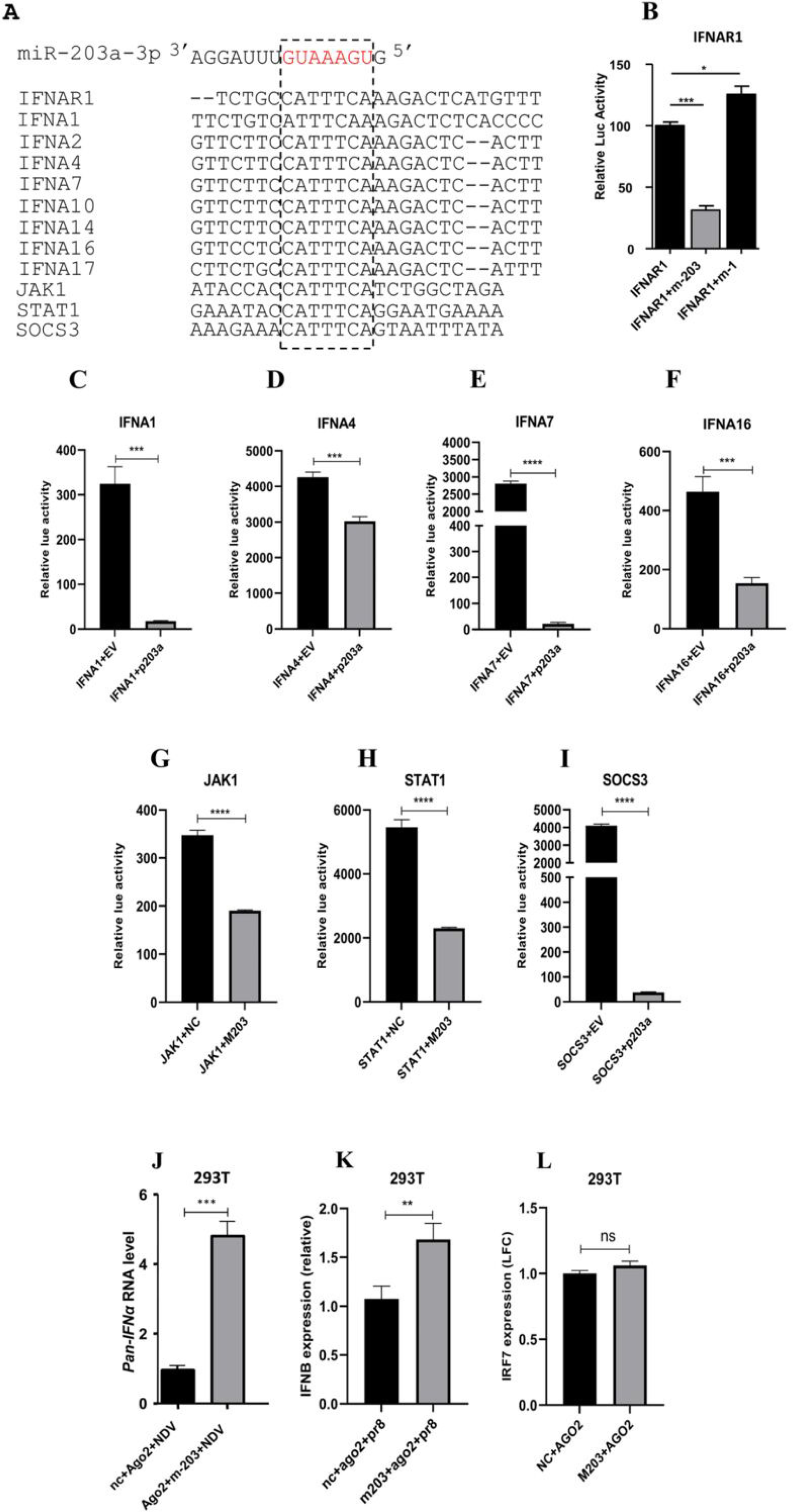
miR-203a-3p(m203) binds with different type-I IFN pathways genes. (A) Showing genes targeted by the m203. (B-I) Type-I IFN pathway genes IFNAR1, IFNA1, IFNA4, IFNA7, IFNA16, JAK1, STAT1 UTRs were cloned into the pMIR-Report vector and transfected with p203 or empty vector, H293T cells were transfected with FNAR1-Luc, IFNA1-Luc, IFNA4-Luc, IFNA7-Luc, IFNA16-Luc, I, JAK1-Luc, STAT1-Luc, SOCS3-Luc,50 ng of pRL-TK, 100 ng of full-length wild-type (WT) 3′UTR, along with 350 ng of the plasmid encoding miR-203 or with 20 nM miR-203 mimic, after 48 hr. cells were lysed and subjected to luciferase assay. (J-K) to confirm the binding of miR-203a-3p to Type-I IFN we transfected the 293T cells with 100 nM miR-203 mimic or mimic negative control (NC) after 24 hours did NDV or PR8 infection at an MOI of 1.5 and samples were collected at 24 hours. RNA was measured by qRT-PCR. The level of significance is indicated by * (p < 0.01), ** (p < 0.001) *** (p < 0.0001), **** (p < 0.00001).

### Viral load upon overexpression of miR-203a-3p

In order to investigate the physiological role of miR-203a-3p during virus replication, A549 cells were transfected with p203, while the empty vector served as control. Subsequently, the transfected cells were infected with Newcastle Disease Virus (NDV) or A/PR8/H1N1 at a multiplicity of infection (MOI) of 1.5. We then quantified the viral load at early (6, 24 hours) and late (48 hours) time points to assess the impact of miR-203a-3p on viral replication. We measured the expression profile of PR8 PB1 gene at 12,24 and 48 hours and the NDV L-pol gene expression which encodes the large polymerase of NDV (referred to as NDV RNA) at 6,12 and 48 hours. Our findings revealed a significant increase in the A/PR8/H1N1 and NDV RNA upon transfection with p203, as compared to the control (Figure 3A-C). The result for NDV was further validated by performing FACS analysis with miR-203a-3p mimic and anti-miR-203a-3p, in presence of mimic viral load increased whereas in presence of anti-miR203a-3p NDV load was found to be reduced (fig. 3 D-E).

**Figure 3:**
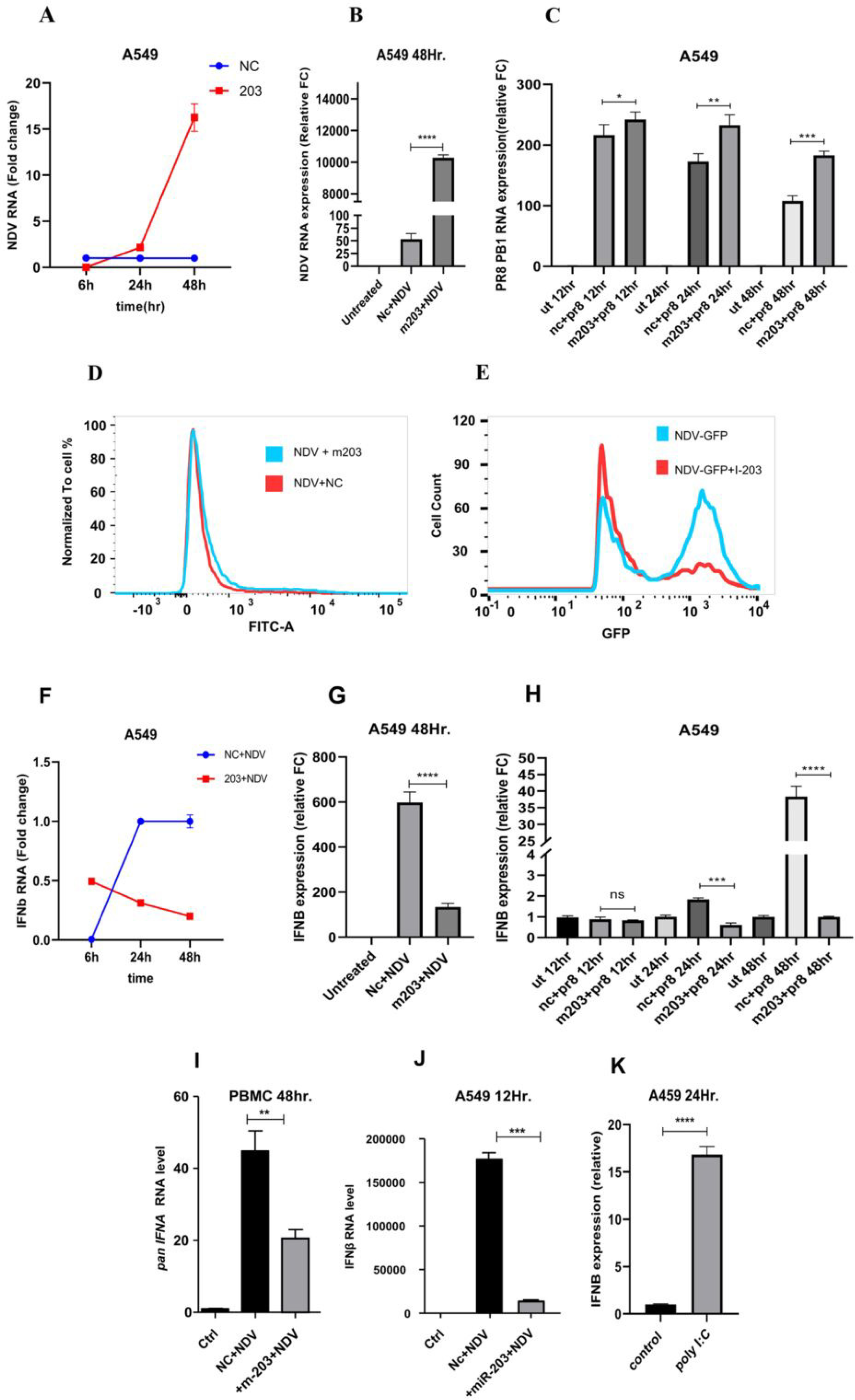
Over expression of miR-203a-3p enhance viral replication: (A-C) A549 transfected with m203a(miR-203a-3p) or NC (negative control) at 50 nM, followed by infection with NDV and PR8 at an MOI of 1.5, samples were collected at indicated time points. Relative expression of L-pol. of NDV (A, B), PB1 gene of PR8 (C, D), RNA was measured by qRT-PCR. (D) to confirm viral load through Fluorescence-activated cell sorting (FACS), A549 cells transfected with NC or m203 at 50 nM followed by NDV infection at an MOI of 1.5, after 24 hr. cells were lysed with trypsin and washed with 1x PBS fixed with formaldehyde (4%) and subjected to FITC-A analysis. (E) A549 cells were transfected with NC or m203 inhibitor after 24hr. Cells were lysed with trypsin and washed with 1x PBS and fixed with formaldehyde (4%) and subjected to FITC-A analysis. (F-H) A549 transfected with m203a(miR-203a-3p) or NC (negative control) at 50 nM, followed by infection with NDV and PR8 at an MOI of 1.5, samples were collected at indicated time points and IFNB expression was measured by qRT-PCR. The level of significance is indicated by * (p < 0.01), ** (p < 0.001) *** (p < 0.0001), **** (p < 0.00001).

**Figure 4:**
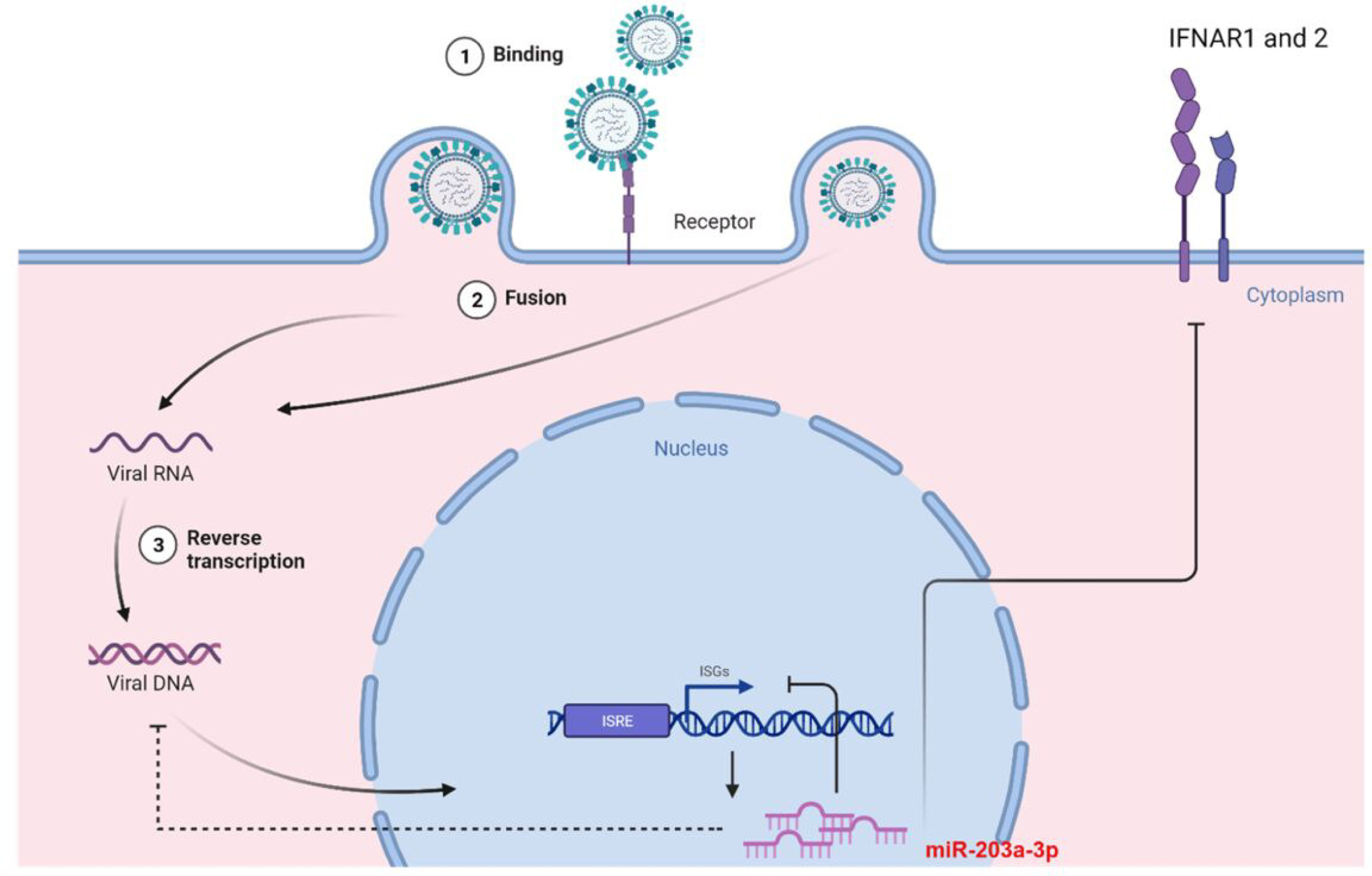
This graphical model illustrates the upregulation of miR-203a-3p expression upon A/PR8/H1N1 and NDV infection. Subsequently, miR-203a-3p binds to the 3’UTR of type-I IFN pathway and ISGs transcripts, and downregulates their expression which in turn facilitates the viral replication. Notably, miR-203a-3p exhibits a weak binding affinity to the viral genome, but its effect on viral replication appears insignificant.

### miR-203a-3p dependent suppression of type-I IFN

To enhance the validation of the impact of miR-203a-3p on type-I IFN expression, we employed a dual approach. Firstly, we transfected A549 cells with a miR-203a-3p mimic. Additionally, we subjected cells to infection with A/PR8/H1N1 or NDV. The virus infection further enhances the miR-203a-3p expression in turn it suppress its targets with mimic. This approach demonstrated a notable decrease in IFN-β, transcript levels across various time points during A/PR8/H1N1 and NDV viral infections (fig. 3F-H), we also checked the expression of pan-α at later time point (fig. 3I). Moreover, to ensure specificity, we solely assessed IFN-β transcript levels through transfection of the poly I:C (see Fig. 3K).

## Discussion

The innate immune system relies on various sensors to detect viral components and initiate an immune response. These sensors include pattern recognition receptors (PRRs) such as Toll-like receptors (TLRs), retinoic acid-inducible gene I (RIG-I)-like receptors (RLRs), nucleotide-binding oligomerization domain (NOD)-like receptors (NLRs), and cytosolic DNA sensors(41, 42). Influenza virus triggers multiple signaling pathways, causing changes in immune and non-immune mediators and regulatory noncoding (small and long) RNAs. These alterations contribute to the immune response and viral clearance(43, 44). Understanding these processes is important for developing effective antiviral strategies. In our study, we found such a microRNA miR-203a-3p. We analyze the publicly available GEO database to identify specific prominently upregulated miRNAs in response to H7N1, HCV, and HIV+HPV virus infections. All data sets show increased expression of miR-203a-3p. To understand the functional implications of miR-203a-3p in the context of viral infections, we performed pathway analysis of microRNA-targeted genes using Enrichr developed by the Ma’ayan Lab. intriguingly, the analysis revealed that miR-203a-3p specifically targets the 3’UTR of type-I interferon pathway genes. Type-I interferon response is a crucial aspect of the host’s innate immune defense against viral infections. Type-I interferon (IFN-I) functions through complex molecular cascade, This cascade results in the activation of numerous Interferon-Stimulated Genes (ISGs), essential for fighting viral infections, and other wide array of genes particularly inducing apoptosis in virally infected cells and to protect the uninfected cells (45, 46). Recent studies have revealed a similar role for Type-III interferon (IFN-III), especially noticeable in epithelial cells on surfaces the main targets of influenza A virus (IAV). When infected by a virus, these cells initiate a dual response by releasing both IFN-I and IFN-III, a process that relies on the activation of RIG-I/MAVS pathway(47).Virus employies different strategies to escape the host’s immune system like cytokine escape, antigenic change, antigen presentation and evasion of natural killer cells(48, 49). It has recently been established in research articlers that the pathogenic insult is also associated with the cellular miRNA expression, and altered miRNA expression leads to the miRNA-mediated silencing of host and viral transcripts(50–52). In contrast of miRNA mediated host mRNA Transcript silencing, here in our study we found that the expression of miR-203a-3p was significantly elevated upon infection with NDV and A/PR8/H1N1 as well as upon transfection with polyinosinic-polycytidylic acid (I: C)], a synthetic analog of viral double-stranded RNA that mimics viral infection.

We observed the upregulated miR-203a-3p promotes the A/PR8/H1N1 and NDV replication via suppressing the type-I IFN pathway. Meanwhile upregulated miR-203a-3p also targets the viral genome but the binding of miR-203a-3p with the viral genome is not efficient to limit the viral replication. The comprehensive analyses strongly indicate that miR-203a-3p plays a pivotal role in modulating the intricate host-virus interaction.Taken together analysis and results shows that A/PR8/H1N1 and NDV find a way to evade the host’s immune system via overexpression of miR-203a-3p. these findings highlight the pivotal role of miR-203a-3p in promoting the A/PR8/H1N1 and NDV virus replication. It accomplishes this by targeting type-I IFN signaling pathway. By promoting the viral replication and modulating the expression of key immune response genes, in this study miR-203a-3p emerges as a crucial player during RNA virus infections. Further exploration of miR-203a-3p’s functional implications and its potential as a therapeutic target could pave the way for novel antiviral strategies.

## Statical analysis

All experiments were meticulously conducted, ensuring the inclusion of appropriate control samples or mock-transfected samples. Each experiment was independently replicated two or three times to ensure robustness and reliability of the results. The data analysis was performed using GraphPad Prism software, a widely recognized tool for statistical analysis. For experiments involving two groups, we employed Student’s two-tailed unpaired t-test, This comprehensive approach guarantees the accuracy and validity of our findings.

## Acknowledgment

We thank R. Fouchier for providing the A/PR8/H1N1 reverse genetics system, P. Palese for green fluorescent protein-expressing NDV (NDV-GFP), and IISER Bhopal for the Central Instrumentation Facility. Generous financial support by the DBT, New Delhi, and SERB, New Delhi is gratefully acknowledged. P.K., thank DBT for awarding SRF.

## Author Contribution

P.K, A.K, Ak.K, and H.K conceptualized the study. P.K and A.K performed experiments.

A.K performed analysis for data available on repository, P.K, A.K experiments on H1N1 and NDV in A549, 293T, HeLa, PBMC. P.K, Ak.K, and H.K prepared the manuscript and H.K supervised the entire project.

---

**Supplementary figure S1:** Pathway analysis was done on miR-203a-3p targeted transcripts using Enrichr. (A) The BioPlanet gene pathway analysis revealed miR-203a-3p targeting the regulation of interferon alpha signaling transcripts, including TRAF6-mediated IRF7 activation. (B) A bar chart displays the top five enriched terms alongside their corresponding p-values. Each bar is color-coded based on its p-value (C) Gene pathways are presented in a table, ranked by P-value and combined score. (D) Uniform Manifold Approximation and Projection (UMAP) visualizes the gene pathways, ranked according to odds ratio and P-value. (E) miR-203a-3p expression was assessed in A/PR8/H1N1 and NDV infections at different MOIs over 24 hours. (F) A549 cells were transfected with poly I:C, and miR203a-3p expression was observed via RT-PCR after 24 hours. (G) A549 cells were transfected with either ISRE alone or ISRE combined with miR-203, or with ISRE combined with miRNA negative control (m-1). After 24 hours, the cells were induced with IFN-β.

**Supplementary figure S2:** Pathway analysis was conducted using Enrichr to explore the targeted transcripts of miR-378c and miR-199a-5p. (A-B) The top 500 miR-378c targeted transcripts were used as input in Enrichr, yet no significant pathway related to the immune system was identified. (C-D) Similarly, the same analysis was performed for miR-199a-5p, but no immune-related pathways were found in this case as well.

## REFERENCES

1. Akira S, Uematsu S, Takeuchi O. 2006. Pathogen recognition and innate immunity. Cell 124:783–801.

2. Takeuchi O, Akira S. 2009. Innate immunity to virus infection. Immunol Rev 227:75–86.

3. Katze MG, He Y, Gale M. 2002. Viruses and interferon: a fight for supremacy. Nat Rev Immunol 2:675–687.

4. Clarke C, Henry M, Doolan P, Kelly S, Aherne S, Sanchez N, Kelly P, Kinsella P, Breen L, Madden SF, Zhang L, Leonard M, Clynes M, Meleady P, Barron N. 2012. Integrated miRNA, mRNA and protein expression analysis reveals the role of post-transcriptional regulation in controlling CHO cell growth rate. BMC Genomics 13:656.

5. Hedl M, Zheng S, Abraham C. 2014. The *IL18RAP* Region Disease Polymorphism Decreases IL-18RAP/IL-18R1/IL-1R1 Expression and Signaling through Innate Receptor–Initiated Pathways. J Immunol 192:5924–5932.

6. Newton K, Dixit VM. 2012. Signaling in innate immunity and inflammation. Cold Spring Harb Perspect Biol 4:a006049.

7. Forti RL, Mitchell WM, Hubbard WC, Workman RJ, Forbes JT. 1984. Pleiotropic activities of human interferons are mediated by multiple response pathways. Proc Natl Acad Sci U S A 81:170–174.

8. Kovarik P, Stoiber D, Novy M, Decker T. 1998. Stat1 combines signals derived from IFN-gamma and LPS receptors during macrophage activation. EMBO J 17:3660–3668.

9. Köster M, Hauser H. 1999. Dynamic redistribution of STAT1 protein in IFN signaling visualized by GFP fusion proteins. Eur J Biochem 260:137–144.

10. Dussurget O, Bierne H, Cossart P. 2014. The bacterial pathogen Listeria monocytogenes and the interferon family: type I, type II and type III interferons. Front Cell Infect Microbiol 4:50.

11. Chapman EJ, Prokhnevsky AI, Gopinath K, Dolja VV, Carrington JC. 2004. Viral RNA silencing suppressors inhibit the microRNA pathway at an intermediate step. Genes Dev 18:1179–1186.

12. Srivastava R, Daulatabad SV, Srivastava M, Janga SC. 2020. Role of SARS-CoV-2 in Altering the RNA-Binding Protein and miRNA-Directed Post-Transcriptional Regulatory Networks in Humans. Int J Mol Sci 21:7090.

13. Sullivan CS, Ganem D. 2005. MicroRNAs and viral infection. Mol Cell 20:3–7.

14. Jopling CL, Yi M, Lancaster AM, Lemon SM, Sarnow P. 2005. Modulation of hepatitis C virus RNA abundance by a liver-specific MicroRNA. Science 309:1577–1581.

15. Lecellier C-H, Dunoyer P, Arar K, Lehmann-Che J, Eyquem S, Himber C, Saïb A, Voinnet O. 2005. A cellular microRNA mediates antiviral defense in human cells. Science 308:557–560.

16. Messina NL, Clarke CJP, Johnstone RW. 2016. Constitutive IFNα/β signaling maintains expression of signaling intermediaries for efficient cytokine responses. JAK-STAT 5:e1173804.

17. Gough DJ, Messina NL, Hii L, Gould JA, Sabapathy K, Robertson APS, Trapani JA, Levy DE, Hertzog PJ, Clarke CJP, Johnstone RW. 2010. Functional crosstalk between type I and II interferon through the regulated expression of STAT1. PLoS Biol 8:e1000361.

18. Brzózka K, Pfaller C, Conzelmann K-K. 2007. Signal transduction in the type I interferon system and viral countermeasures. Signal Transduct 7:5–19.

19. McNab F, Mayer-Barber K, Sher A, Wack A, O’Garra A. 2015. Type I interferons in infectious disease. Nat Rev Immunol 15:87–103.

20. de Weerd NA, Nguyen T. 2012. The interferons and their receptors--distribution and regulation. Immunol Cell Biol 90:483–491.

21. Hartman SE, Bertone P, Nath AK, Royce TE, Gerstein M, Weissman S, Snyder M. 2005. Global changes in STAT target selection and transcription regulation upon interferon treatments. Genes Dev 19:2953–2968.

22. Michalska A, Blaszczyk K, Wesoly J, Bluyssen HAR. 2018. A Positive Feedback Amplifier Circuit That Regulates Interferon (IFN)-Stimulated Gene Expression and Controls Type I and Type II IFN Responses. Front Immunol 9:1135.

23. Cho SS, Bacon CM, Sudarshan C, Rees RC, Finbloom D, Pine R, O’Shea JJ. 1996. Activation of STAT4 by IL-12 and IFN-alpha: evidence for the involvement of ligand-induced tyrosine and serine phosphorylation. J Immunol Baltim Md 1950 157:4781–4789.

24. Nguyen KB, Watford WT, Salomon R, Hofmann SR, Pien GC, Morinobu A, Gadina M, O’Shea JJ, Biron CA. 2002. Critical role for STAT4 activation by type 1 interferons in the interferon-gamma response to viral infection. Science 297:2063–2066.

25. Ingle H, Kumar S, Raut AA, Mishra A, Kulkarni DD, Kameyama T, Takaoka A, Akira S, Kumar H. 2015. The microRNA miR-485 targets host and influenza virus transcripts to regulate antiviral immunity and restrict viral replication. Sci Signal 8:ra126.

26. Clough E, Barrett T. 2016. The Gene Expression Omnibus Database. Methods Mol Biol Clifton NJ 1418:93–110.

27. Agarwal V, Bell GW, Nam J-W, Bartel DP. 2015. Predicting effective microRNA target sites in mammalian mRNAs. eLife 4:e05005.

28. Enright AJ, John B, Gaul U, Tuschl T, Sander C, Marks DS. 2003. MicroRNA targets in Drosophila. Genome Biol 5:R1.

29. Chen Y, Wang X. 2020. miRDB: an online database for prediction of functional microRNA targets. Nucleic Acids Res 48:D127–D131.

30. Quillet A, Saad C, Ferry G, Anouar Y, Vergne N, Lecroq T, Dubessy C. 2019. Improving Bioinformatics Prediction of microRNA Targets by Ranks Aggregation. Front Genet 10:1330.

31. Quillet A, Anouar Y, Lecroq T, Dubessy C. 2021. Prediction methods for microRNA targets in bilaterian animals: Toward a better understanding by biologists. Comput Struct Biotechnol J 19:5811–5825.

32. Gagliardi M, Matarazzo MR. 2016. RIP: RNA Immunoprecipitation. Methods Mol Biol Clifton NJ 1480:73–86.

33. Meister G, Landthaler M, Patkaniowska A, Dorsett Y, Teng G, Tuschl T. 2004. Human Argonaute2 mediates RNA cleavage targeted by miRNAs and siRNAs. Mol Cell 15:185–197.

34. Mendoza EJ, Manguiat K, Wood H, Drebot M. 2020. Two Detailed Plaque Assay Protocols for the Quantification of Infectious SARS-CoV-2. Curr Protoc Microbiol 57:ecpmc105.

35. Wulff NH, Tzatzaris M, Young PJ. 2012. Monte Carlo simulation of the Spearman-Kaerber TCID50. J Clin Bioinforma 2:5.

36. Kanehisa M, Goto S. 2000. KEGG: kyoto encyclopedia of genes and genomes. Nucleic Acids Res 28:27–30.

37. Kanehisa M, Furumichi M, Sato Y, Kawashima M, Ishiguro-Watanabe M. 2023. KEGG for taxonomy-based analysis of pathways and genomes. Nucleic Acids Res 51:D587–D592.

38. Chen EY, Tan CM, Kou Y, Duan Q, Wang Z, Meirelles GV, Clark NR, Ma’ayan A. 2013. Enrichr: interactive and collaborative HTML5 gene list enrichment analysis tool. BMC Bioinformatics 14:128.

39. Kuleshov MV, Jones MR, Rouillard AD, Fernandez NF, Duan Q, Wang Z, Koplev S, Jenkins SL, Jagodnik KM, Lachmann A, McDermott MG, Monteiro CD, Gundersen GW, Ma’ayan A. 2016. Enrichr: a comprehensive gene set enrichment analysis web server 2016 update. Nucleic Acids Res 44:W90–97.

40. Ikeda K, Satoh M, Pauley KM, Fritzler MJ, Reeves WH, Chan EKL. 2006. Detection of the Argonaute Protein Ago2 and microRNAs in the RNA Induced Silencing Complex (RISC) Using a Monoclonal Antibody. J Immunol Methods 317:38–44.

41. Li D, Wu M. 2021. Pattern recognition receptors in health and diseases. Signal Transduct Target Ther 6:291.

42. Takaoka A, Wang Z, Choi MK, Yanai H, Negishi H, Ban T, Lu Y, Miyagishi M, Kodama T, Honda K, Ohba Y, Taniguchi T. 2007. DAI (DLM-1/ZBP1) is a cytosolic DNA sensor and an activator of innate immune response. Nature 448:501–505.

43. Kasuga Y, Zhu B, Jang K-J, Yoo J-S. 2021. Innate immune sensing of coronavirus and viral evasion strategies. Exp Mol Med 53:723–736.

44. Kawai T, Akira S. 2006. Innate immune recognition of viral infection. Nat Immunol 7:131–137.

45. Barber GN. 2001. Host defense, viruses and apoptosis. Cell Death Differ 8:113–126.

46. Nukui M, Mori Y, Murphy EA. 2015. A human herpesvirus 6A-encoded microRNA: role in viral lytic replication. J Virol 89:2615–2627.

47. Crotta S, Davidson S, Mahlakoiv T, Desmet CJ, Buckwalter MR, Albert ML, Staeheli P, Wack A. 2013. Type I and type III interferons drive redundant amplification loops to induce a transcriptional signature in influenza-infected airway epithelia. PLoS Pathog 9:e1003773.

48. LUCAS M, KARRER U, LUCAS A, KLENERMAN P. 2001. Viral escape mechanisms – escapology taught by viruses. Int J Exp Pathol 82:269–286.

49. Rubio-Casillas A, Redwan EM, Uversky VN. 2022. SARS-CoV-2: A Master of Immune Evasion. Biomedicines 10:1339.

50. Sharma S, Chatterjee A, Kumar P, Lal S, Kondabagil K. 2020. Upregulation of miR-101 during Influenza A Virus Infection Abrogates Viral Life Cycle by Targeting mTOR Pathway. Viruses 12:444.

51. Othumpangat S, Beezhold DH, Umbright CM, Noti JD. 2021. Influenza Virus-Induced Novel miRNAs Regulate the STAT Pathway. Viruses 13:967.

52. Huang S-Y, Huang C-H, Chen C-J, Chen T-W, Lin C-Y, Lin Y-T, Kuo S-M, Huang C-G, Lee L-A, Chen Y-H, Chen M-F, Kuo R-L, Shih S-R. 2019. Novel Role for miR-1290 in Host Species Specificity of Influenza A Virus. Mol Ther -Nucleic Acids 17:10–23.

